# Oviduct fluid metabolic regulation of embryonic genome methylation

**DOI:** 10.1101/2025.06.13.659599

**Authors:** Dailin M. Fuego, Iebu Devkota, Zachary L. Bonomo, Yuxia Li, Xujia Zhang, Mahmoud F. Dondeti, Fabrizio Donnarumma, Antonios Matsakas, Amanda L. Patterson, Cathleen C. Williams, Anastasios Vourekas, Xing Fu, Kenneth R. Bondioli, Constantine A. Simintiras

## Abstract

Adverse maternal health and lifestyle during pregnancy can program offspring susceptibility to noncommunicable diseases in adulthood. Oviduct fluid facilitates key preimplantation milestones, including embryonic genome activation. However, the regulatory mechanisms governing oviduct fluid composition, and the developmental consequences of its disruption, remain unclear. Capitalizing on conserved dynamics between human and bovine embryonic genome activation, we first established and characterized bovine oviduct epithelial organoids as a model system. Organoids were then exposed to Δ⁹-tetrahydrocannabinol (**THC**) and cannabidiol (**CBD**) to test the hypothesis that exocannabinoids alter intra-organoid fluid composition.

Treatment altered the organoid transcriptome and intra-organoid fluid metabolome, notably elevating 5′-deoxyadenosine levels. Subsequent embryo culture with 5′-deoxyadenosine during embryonic genome activation resulted in aberrant DNA methylation patterning – independently of direct THC and CBD exposure. These findings identify a novel, indirect, mechanism by which maternal exposures disrupt embryonic development through changes in oviductal metabolite secretions.

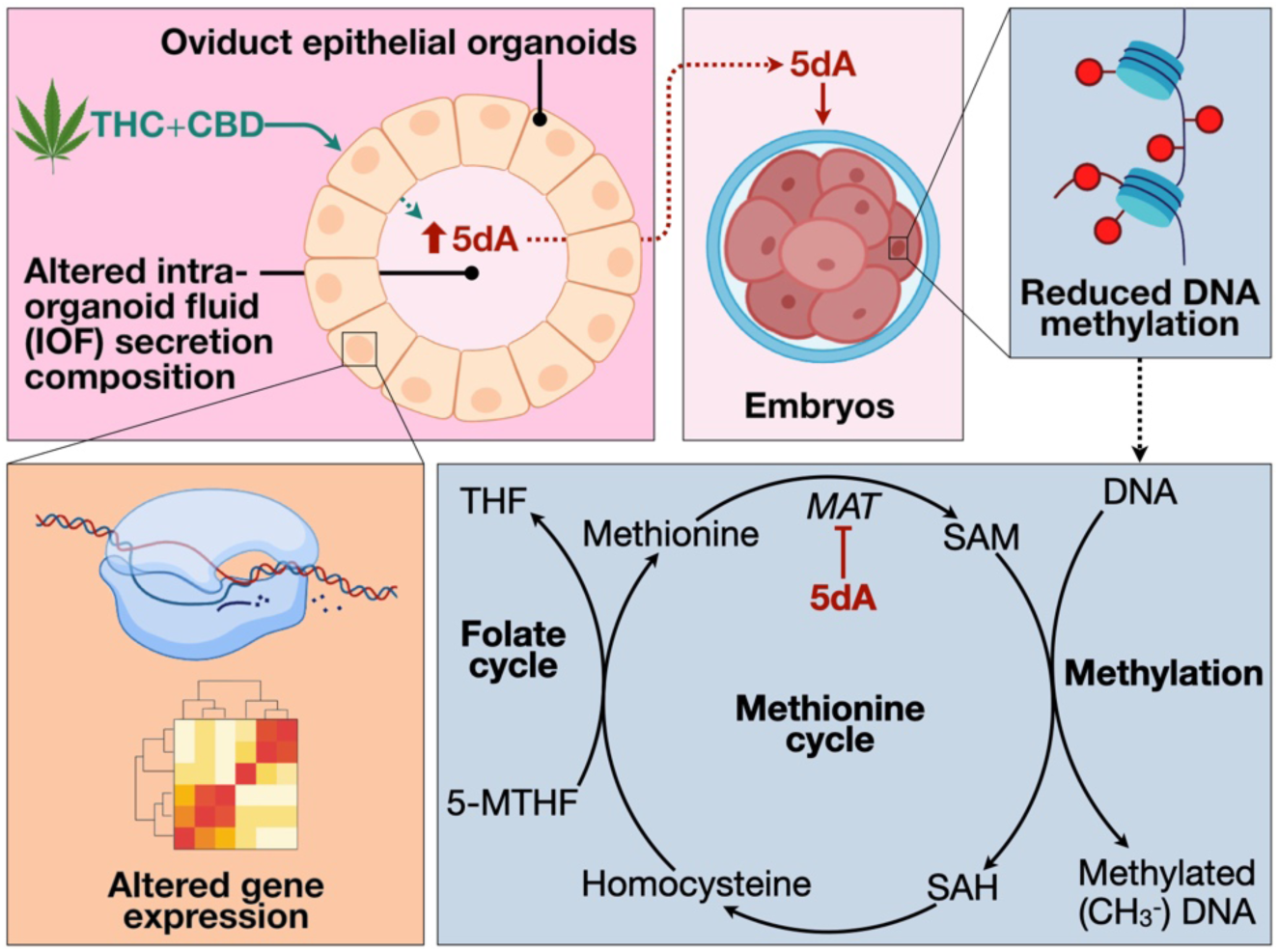
Graphical Abstract.

## INTRODUCTION

Oviduct fluid facilitates critical early reproductive events including fertilization, early cleavage, and embryonic genome activation. Occurring between Days 2-5 post-fertilization, embryonic genome activation marks the embryonic transition from maternal transcript reliance to autonomous gene expression^1^, driven by extensive epigenetic reprogramming – notably genomic DNA re-methylation^2^. This epigenetic plasticity renders the embryo acutely sensitive to its microenvironment, with perturbations capable of negatively influencing its developmental trajectory, in line with the developmental origins of health and disease (**DOHaD**) paradigm^3^. The DOHaD framework states that prenatal exposures shape susceptibility to noncommunicable diseases, including cardiometabolic disorders^4^, atopic conditions^5^, malignancies^6^, and neurodevelopmental impairment^7^, among other ailments^8^. While the impact of developmental programming spans gestation and early life, mounting evidence from human and animal models underscores the peri-conceptional period as a critical window of vulnerability and opportunity^9^.

One of the clearest demonstrations of human periconceptual sensitivity comes from assisted reproductive technologies, such as intra-cytoplasmic sperm injection and *in vitro* fertilization – which entirely bypass the oviduct. Offspring conceived via such assisted reproductive technologies show an increased incidence of disorders linked to aberrant DNA methylation^10^. This phenomenon is partly attributable to the plasticware used to handle and culture embryos *in vitro*^11^. However, naturally conceived embryos are also susceptible to DOHaD-associated phenotypes. For example, maternal prenatal cannabis use is linked to increased risks of autism spectrum disorder, intellectual disability, and other neurobehavioral impairments in offspring^12^. These findings are particularly concerning since cannabis is the most consumed illicit substance in the United States, with prenatal use increasing over time, and peaking during the first trimester compared to the second and third^13^.

Dominant metabolites in circulation following cannabis use are Δ⁹-tetrahydrocannabinol (**THC**) and cannabidiol (**CBD**)^14^, while the two primary cannabinoid receptors are CB1 and CB2^15^. CB1 is expressed throughout the human oviduct^16^, whereas CB2 expression has not been detected to date. Similarly, the mouse oviduct expresses CB1, but not CB2^17^. In contrast, both CB1 and CB2 are expressed in the bovine oviduct^18^. Data on cannabinoid receptor expression in human preimplantation embryos remain limited; however, both CB1 and CB2 are expressed in the human oocyte^19^ and mouse preimplantation embryo^20^. In contrast, the bovine embryo expresses RNA transcripts for CB2 (*CNR2* gene in bovine) but not CB1 (*CNR2*)^21–23^.

Endogenous and exogenous cannabinoid, as well as cannabinoid receptor agonist, supplementation impairs preimplantation mouse embryo development *in vitro*^24^. However, the physiological relevance of these findings is unclear, as it is unknown whether THC and CBD – uncharged, hydrophobic molecules – can transverse the oviductal epithelium to reach the lumen, thereby affecting embryo development directly^25^. Moreover, the existence of a cannabinoid transporter has not been conclusively demonstrated^26–29^. Genetic knockout mice have shown that CB1 deletion increases embryo retention within the oviduct and consequently pregnancy failure^17^, while dual CB1 and CB2 knockout mice exhibit impaired uterine implantation^30^. These findings affirm a critical role for endocannabinoid signaling in pregnancy establishment^31^.

However, a fundamental and unresolved question remains – whether circulating exocannabinoids can disrupt early embryo development indirectly, by modulating oviductal physiology and behavior, rather than, or in addition to, acting on the embryo directly.

Given the ethical and technical constraints surrounding *in vivo* and *in vitro* human oviduct research, the bovine was selected as a model due to the closer alignment with human embryo development than the mouse – particularly in the timing and regulation of embryonic genome activation and epigenetic reprogramming^32,33^. In parallel, organoids are a powerful platform for investigating interstitial fluid composition regulation under controlled *in vitro* conditions. For example, snake venom gland organoids produce biologically active venom within their intra-organoid fluid^34^. Similarly, human lacrimal gland organoids produce tears (*crying* organoids)^35^, human choroid plexus organoids produce cerebrospinal fluid^36^, and human endometrial organoids produce a uterine-like fluid^37^. Moreover, bovine oviduct epithelial organoids have been created and shown to exhibit a transcriptomic response to heat stress^38^.

Here we establish bovine oviduct epithelial organoids as a hormonally responsive *in vitro* model of the oviduct, recapitulating both transcriptomic and intra-organoid fluid profiles. We then demonstrate that THC and CBD exposure alters intra-organoid fluid composition, supporting our hypothesis and revealing downstream effects on embryonic epigenetic programming. These findings offer mechanistic insight into how maternally circulating, DOHaD-relevant xenobiotics, can indirectly impair early embryonic development by disrupting oviduct epithelial behavior.

## RESULTS

### Oviduct organoid establishment and characterization

Our first objective was to establish and characterize bovine oviduct epithelial organoids. Oviducts were collected from estrous synchronized crossbred beef cattle (n=4) euthanized on Day 5 of the estrous cycle – a timepoint coinciding with embryonic genome activation (**Fig**. **1A**). Oviducts ipsilateral to the *corpus luteum* were excised and a piece of the isthmus was dissected for immunofluorescence imaging. Thereafter, the remaining oviduct was flushed with phosphate-buffered saline (**PBS**) to obtain native oviduct fluid, which served as the *in vivo* reference for downstream intra-organoid fluid comparisons. Cells were subsequently obtained from the remaining oviduct and partitioned for either transcriptomic profiling (RNA-seq) or organoid derivation (**Fig**. **1B**). *In vivo* tissue immunofluorescence imaging confirmed epithelial expression of cytokeratin, oviduct-specific glycoprotein 1 (**OVGP1**), and steroid hormone receptors estrogen receptor alpha (**ER⍺**), estrogen receptor beta (**ERβ**), and the progesterone receptor (**PGR**) (**Fig**. **1C**), as expected.

**Figure 1.**
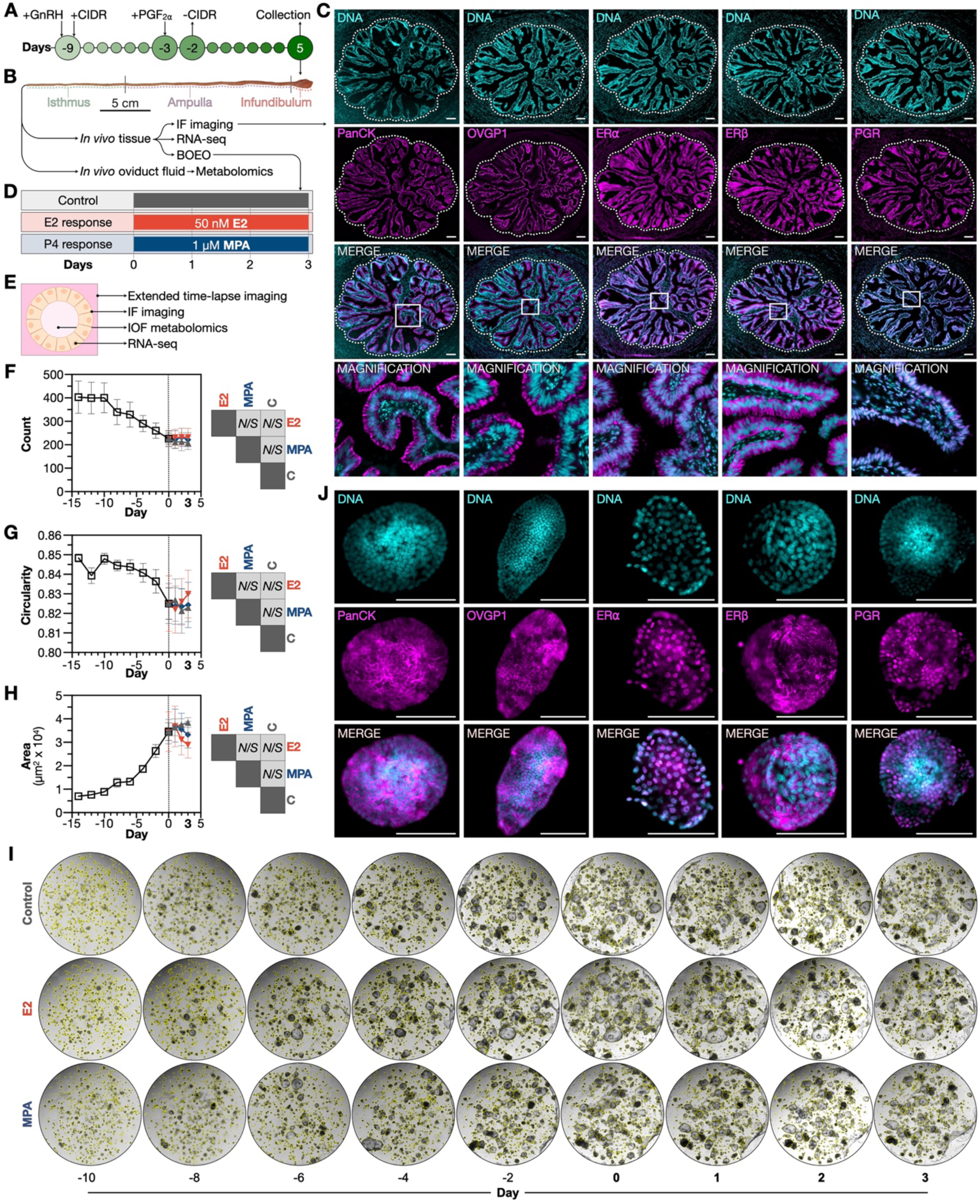
Oviduct organoid characterization. (**A**) Estrous synchronization protocol, including gonadotropin-releasing hormone (GnRH), controlled internal drug release (CIDR) progesterone (P4) insert, and prostaglandin F2⍺ (PGF2⍺) administration, with sample collection on Day 5 post-estrus. (**B**) Sample collection workflow – a portion of the oviduct was retained for immunofluorescence (IF) imaging before whole oviduct fluid and cell collection. (**C**) Representative immunofluorescence images of Day 5 *in vivo* oviduct isthmus histoarchitecture, showing cytokeratin (PanCK), oviduct-specific glycoprotein 1 (OVGP1), estrogen receptors alpha (ER⍺) and beta (ERβ), and P4 receptor (PGR) expression. Dotted lines denote oviduct mucosa and muscularis boundaries. (**D**) Dependent experimental variables – bovine oviduct epithelial organoids (BOEO) were treated with 17β-estradiol (E2), medroxyprogesterone acetate (MPA), or not (control) for 72 h. (**E**) Independent experimental variables, including intra-organoid fluid (IOF) metabolomic profiling. (**F**-**H**) Mean (±SEM) organoid (n=4) morphometric kinetics – (**F**) count, (**G**) circularity, and (**H**) area. No treatment effects observed [not significant (*N*/*S*)]. (**I**) Corresponding representative organoid brightfield images across treatments and time. Note: Yellow borders are software generated. (**J**) Representative organoid immunohistochemistry following E2 treatment. All scale bars: 100 µm.

Following initial derivation and culture, organoids were cryopreserved for three months before regeneration, demonstrating their capacity for biobanking, similarly to other organoids^39^. All experiments were conducted at this first subculture. Specifically, organoids were cultured and monitored by brightfield microscopy every 48 h during a 14-day establishment phase, and every 24 h during subsequent treatment (**Fig**. **1D**). Specifically, to evaluate hormone responsiveness, organoids were treated for 72 h with 17β-estradiol (**E2**) or medroxyprogesterone acetate (**MPA**), a synthetic progestin^40^, at concentrations guided by human endometrial organoid literature^41^.

Post-treatment, one-third of organoids were fixed for immunofluorescence imaging. The remaining were processed for intra-organoid fluid metabolomic profiling and organoid transcriptomic profiling (**Fig**. **1E**). All samples (*in vivo* tissue, organoids, and intra-organoid fluid) were derived from the same animals, enabling direct intra-subject comparisons. Organoid morphology, in terms of count (**Fig**. **1F**), circularity (**Fig**. **1G**), and cross-sectional area (**Fig**. **1H**), did not differ between treatments. Representative images are provided in **Figure 1I**. Immunofluorescence imaging confirmed that organoids maintained an epithelial phenotype, expressing the characteristic OVGP1 and all three steroid receptors, recapitulating *in vivo* expression patterns (**Fig**. **1J**). No marker expression differences were observed between groups.

### Oviduct organoid transcriptomic responsiveness to hormonal stimulation

Despite the absence of morphological changes, RNA-seq revealed robust transcriptional responsiveness to hormonal supplementation. Following quality control (including one MPA-treated replicate excluded due to low read count), principal component analysis (**PCA**) demonstrated partial overlap, but substantial separation, between E2 *vs*. MPA treated groups (**Fig**. **2A**). However, *in vivo* and *in vitro* samples clustered distinctly, likely reflecting cellular heterogeneity (*e*.*g*., stromal and immune cells) absent in epithelial-only organoids. Hierarchical clustering of top differentially expressed genes (**DEG**) identified six distinct functional modules (**Fig**. **2B**), with gene ontology analysis revealing enrichment for ciliation (cluster 1), tRNA aminoacylation (cluster 2), small molecule metabolism (cluster 3), axonemal dynein complex assembly (cluster 4), fertilization (cluster 5), and transmembrane transport (cluster 6), among others (**Fig**. **2C**).

**Figure 2.**
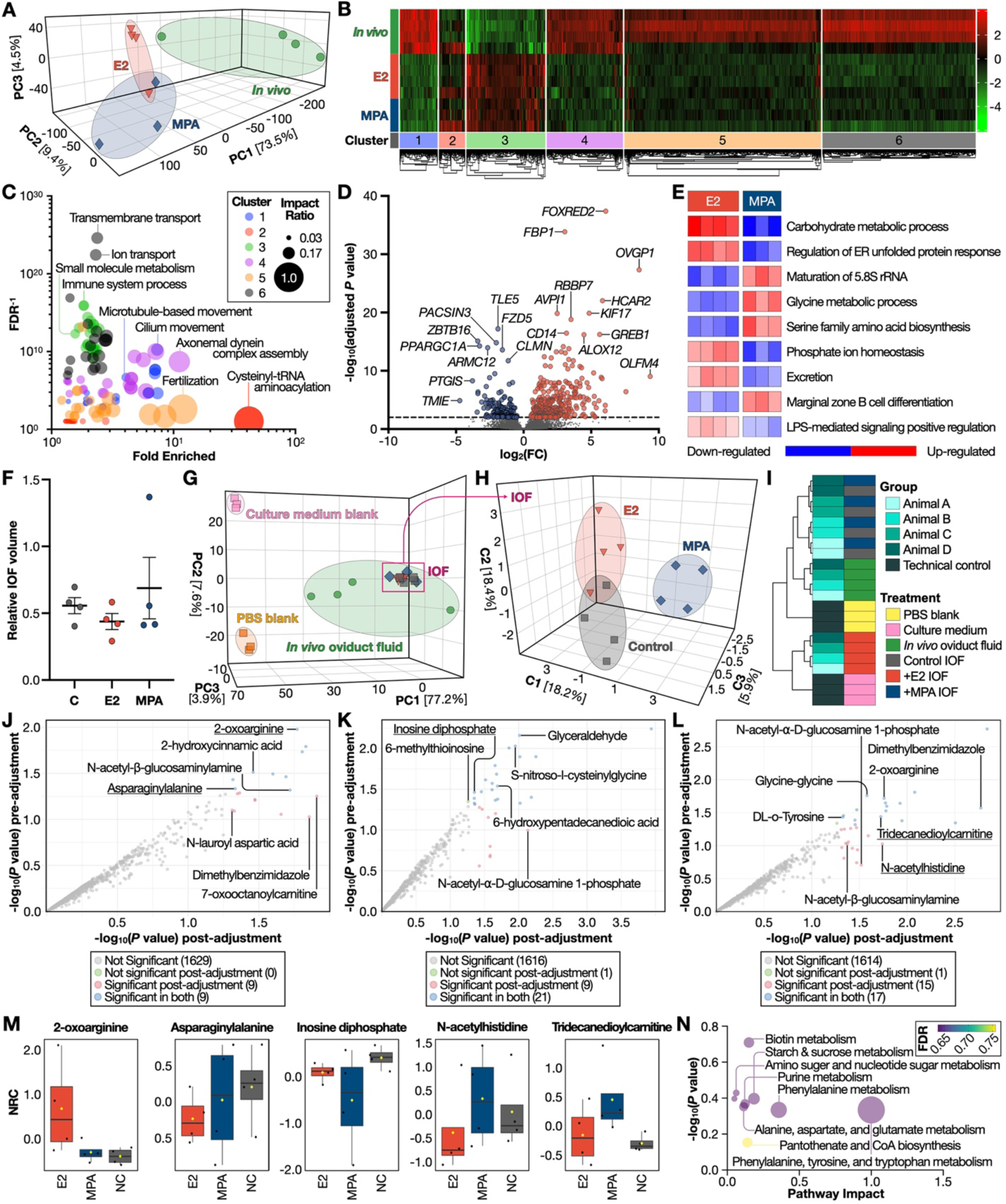
Oviduct organoids are hormonally responsive. (**A**) Principal component (PC) analysis (PCA) of transcriptomic profiles of *in vivo* oviduct tissue (n=4) against 17β-estradiol (E2; n=4) and medroxyprogesterone acetate (MPA; n=3) treated organoids. (**B**) Heatmap of differentially expressed genes (DEG) by treatment, grouped into six clusters. (**C**) Multivariate plot of DEG pathway significance and impact according to cluster. Bubble size is proportional to the impact ratio (number of genes divided by total pathway size), whereas the vertical axis unit is inverse false discovery rate (FDR). (**D**) DEG between organoids treated with E2 *vs*. MPA. Red and blue indicate E2 and MPA upregulation, respectively. (**E**) Parametric gene set enrichment analysis using gene ontology resource *biological process* pathway annotations, highlighting activated (red) and suppressed (blue) processes in E2 *vs*. MPA treated organoids. (**F**) Mean (±SD) intra-organoid fluid (IOF) relative volumes (n=4). No treatment effects observed [not significant (*N*/*S*)]. (**G**) PCA of metabolomic profiles from media and PBS blanks, *in vivo* oviduct fluid, and IOF. (**H**) Sparse partial least squares discriminant analysis (sPLS-DA) of IOF metabolomes across control, E2, and MPA treated organoids. (**I**) IOF metabolomic profile metadata heatmap with hierarchical clustering by animal and treatment. (**J**-**L**) Linear models with covariate (group) adjustment comparing IOF metabolomic profiles – (**J**) control *vs*. E2, (**K**) control *vs*. MPA, and (**L**) E2 *vs*. MPA. (**M**) Boxplots of select metabolite normalized relative concentrations (NRC) across treatments. Central horizontal lines represent the median value with outer box boundaries depicting upper and lower quartile limits. Error bars depict the minimum and maximum distributions, with yellow rhombi representing the mean values. (**N**) Pathway impact analysis using differentially abundant metabolites between E2 *vs*. MPA treated IOF. Bubble size is proportional to pathway impact whereas bubble color is proportional to the FDR.

More specifically, E2 *vs*. MPA supplementation resulted in 876 DEG, with 468 upregulated in the E2 group and 408 upregulated in the MPA treated groups. Notably, E2 upregulated several canonical estrogen-responsive genes, including *OVGP1* and *OLMF4*^42^, while MPA upregulated many progesterone-responsive targets such as *CLMN* and *ZBTB16*^43^ (**Fig**. **2D**). Moreover, parametric gene set enrichment analysis identified multiple differentially regulated metabolic pathways in response to E2 *vs*. MPA (**Fig**. **2E**), further highlighting bovine oviduct epithelial organoid functional divergence in response to E2 *vs*. MPA, despite similar morphology.

### Oviduct organoid secretions mirror in vivo oviduct fluid

We next examined the intra-organoid fluid metabolomic composition from E2 *vs*. MPA treated and control organoids. Total fluid volumes (normalized to total RNA) were similar between groups (**Fig**. **2F**). However, PCA confirmed that intra-organoid fluid profiles clustered separately from both the culture medium and PBS (intra-organoid fluid diluent) controls. Moreover, the intra-organoid fluid metabolome overlapped substantially with native oviduct fluid – further supporting the physiological fidelity of this model. Sparse partial least squares discriminant analysis (**sPLS**-**DA**) further distinguished E2 *vs*. MPA intra-organoid fluid profiles (**Fig**. **2H**), suggesting treatment-specific metabolic shifts, despite the overall similarity between groups observed by PCA (not shown).

Hierarchical clustering analysis revealed treatment as the dominant source of variance, although animal-specific effects were also observed (**Fig**. **2I**). Linear covariate adjustment accounting for this batch effect identified 18, 30, and 32 differentially abundant metabolites (**DAM**) in control *vs*. E2 (**Fig**. **2J**), control *vs*. MPA (**Fig**. **2K**), and E2 *vs*. MPA (**Fig**. **2L**) comparisons, respectively. Select DAM are provided in **Figure 2M**. Moreover, pathway enrichment analysis highlighted pronounced differences in phenylalanine, tyrosine, and tryptophan metabolism, among other pathways, between intra-organoid fluids profiled from E2 *vs*. MPA treated organoids (**Fig**. **2N**).

Together, these results demonstrate that bovine oviduct epithelial organoids, while morphologically consistent, respond to steroidal stimuli at the transcriptomic and metabolomic levels, with intra-organoid fluid composition closely mimicking native oviduct fluid.

### Exocannabinoid exposure alters oviduct organoid morphology

To determine the effects of exocannabinoids on oviduct function, we first confirmed CB1 and CB2 receptor expression in both native bovine oviduct tissue (**Fig**. **3A**) and corresponding organoids (**Fig**. **3B**). Organoids were then treated with: (*a*) E2+MPA (physiological mimic), (*b*) E2+MPA+THC+CBD (termed THC+CBD group) to mimic pathological exposure, or (*c*) vehicle control (**Fig**. **3C**). THC and CBD concentrations [6 µM] were based on prior *in vitro* studies investigating the impact of exocannabinoids on bovine semen^44^ and murine embryos^20^.

**Figure 3.**
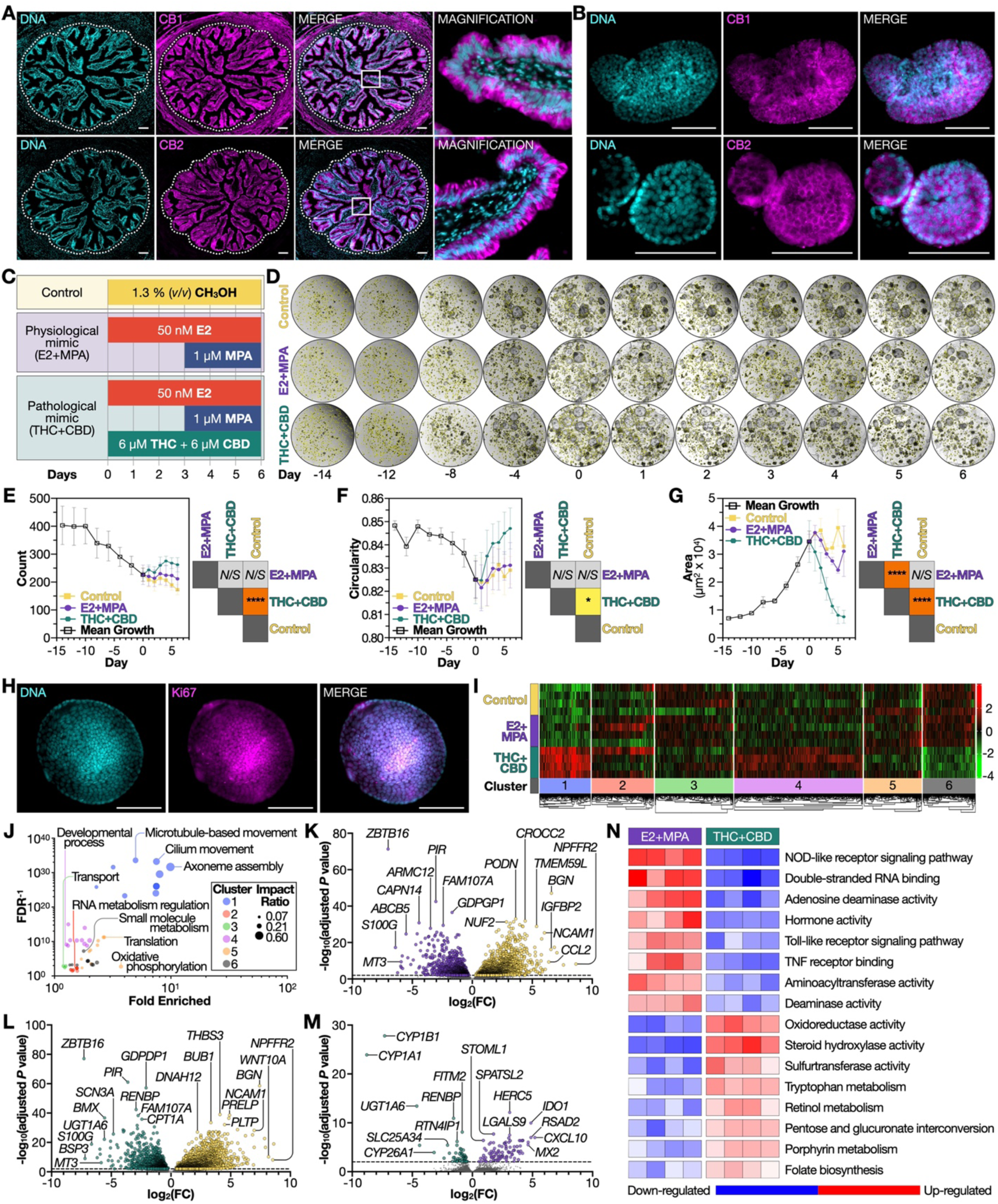
Oviduct organoids respond to pathophysiological stimuli. (**A**) Representative *in vivo* bovine oviduct histoarchitecture at Day 5, showing cannabinoid receptor 1 (CB1) and 2 (CB2) expression. (**B**) Representative control group organoid CB1 and CB2 expression. (**C**) Independent experimental variables – (*i*) 17β-estradiol (E2) with medroxyprogesterone acetate (MPA); (*ii*) E2 and MPA plus Δ⁹-tetrahydrocannabinol (THC) and cannabidiol (CBD) supplementation; or (*iii*) methanol (control) supplementation for 144 h. (**D**) Representative organoid brightfield images across treatments and time. Note: Yellow borders are software generated. (**E**-**G**) Mean (±SEM) organoid (n=4) morphometric kinetics – (**E**) count, (**F**) circularity, and (**G**) area. Select differences observed [*P*≤0.05 (*) and *P*≤0.0001 (****)]. (**H**) Representative organoid Ki67 expression following pathological mimic (THC+CBD) supplementation. (**I**) Heatmap of differentially expressed genes (DEG) by treatment, grouped into six clusters. (**J**) Multivariate plot of DEG pathway significance and impact according to cluster. Bubble size is proportional to the impact ratio (number of genes divided by total pathway size), whereas the vertical axis unit is inverse false discovery rate (FDR). (**K**-**M**) DEG between organoid treatments – (**K**) control *vs*. E2+MPA, (**L**) control *vs*. THC+CBD, and (**M**) E2+MPA *vs*. THC+CBD. Yellow, purple, and teal indicate control, E2+MPA, and THC+CBD gene upregulation, respectively. (**N**) Parametric gene set enrichment analysis using gene ontology resource *molecular function* and Kyoto Encyclopedia of Genes and Genomes (KEGG) pathway annotations, highlighting activated (red) and suppressed (blue) processes in E2+MPA *vs*. THC+CBD treated organoids. All scale bars: 100 µm.

Moreover, treatment duration was extended from 72 h to 144 h to capture sustained transcriptional and secretory changes. Organoids were imaged at regular intervals (**Fig**. **3D**), and morphometric parameters quantified as above. Intriguingly, THC+CBD treatment induced measurable morphological changes – even after 72 h supplementation (not shown). By 144 h, organoid count (**Fig**. **3E**) and circularity (**Fig**. **3F**) increased relative to control, and cross-sectional area decreased relative to both control and E2+MPA (**Fig**. **3G**). Ki67 labelling confirmed this exocannabinoid-induced phenotype was not due to cytotoxicity (**Fig**. **3H**).

### Exocannabinoid exposure alters the oviduct organoid transcriptome

RNA-seq revealed organoid transcriptional remodeling in response to exocannabinoid supplementation for 144 h. DEG clustered into six distinct modules (**Fig**. **3I**), with gene ontology classification highlighting a wide range of impacted pathways, including ciliation (cluster 1), RNA metabolism regulation (cluster 2), transport (cluster 3), developmental processes (cluster 4), translation (cluster 5), and small molecule metabolism (cluster 6), among others (**Fig**. **3J**).

Comparison of E2+MPA *vs*. control organoid transcripts identified 5,009 DEG, including upregulation of *ZBTB16* (PLZF), a direct PGR target and transcription factor essential for murine uterine receptivity^45^. Conversely, *CROCC2*, which modifies cilia dynamics^46^, and is a predicted DEG in the mouse oviduct during pregnancy^47^, was downregulated (**Fig**. **3K**).

DEG between THC+CBD *vs*. control organoid totaled 5,650 and included *GDPD1* (GDE4), which was upregulated and hydrolyzes lysoglycerophospholipids to produce lysophosphatidic acid and free amines^48^, and *FAM107* (DRR1), a stress-inducible actin bundling protein linked to cytoskeletal remodelling^49^. Among the downregulated genes were *BUB1*, a core spindle assembly checkpoint component^50^ also involved in telomere replication^51^, and *THBS3*, whose expression is associated with suppressed oxidative phosphorylation and oncogenesis^52^ (**Fig**. **3L**).

Direct transcript abundance comparison between E2+MPA *vs*. THC+CBD treatments revealed 325 DEG attributable to THC+CBD supplementation. These included a robust induction of cytochrome P450 enzyme genes (*e*.*g*., CYP1B1), consistent with xenobiotic metabolism^53^, and UGT1A6, also involved in cellular detoxification^54^. Meanwhile, downregulated DEG in the same comparison included Type I interferon response inflammation regulators, such as *HERC5*^55^, *IDO1*^56^, *CXCL10*^57^, *RIGI*^58^, *STOML1*^59^, and *RSAD2*^60^, among others (**Fig**. **3M**). Finally, parametric gene set enrichment analysis confirmed and extended these findings, revealing DEG enrichment for pathways related to metabolism and immune responses between E2+MPA *vs*. THC+CBD treated organoids (**Fig**. **3N**).

### Exocannabinoid exposure disrupts oviduct organoid secretion composition

We next profiled the intra-organoid fluid metabolome to evaluate whether disruptions in the organoid transcriptome were reflected in secretory output. Total intra-organoid fluid volumes (normalized to total RNA) remained unchanged across treatments (**Fig**. **4A**). However, PCA revealed clear separation between intra-organoid fluid profiles from E2+MPA *vs*. THC+CBD organoids (**Fig**. **4B**). We also compared intra-organoid fluid metabolomes from this experiment (144 h treatment) to the previous (72 h treatment), finding that time-in-culture also appears to contribute to variance. sPLS-DA supported distinct fluid clustering based on organoid THC+CBD supplementation (**Fig**. **4C**).

**Figure 4.**
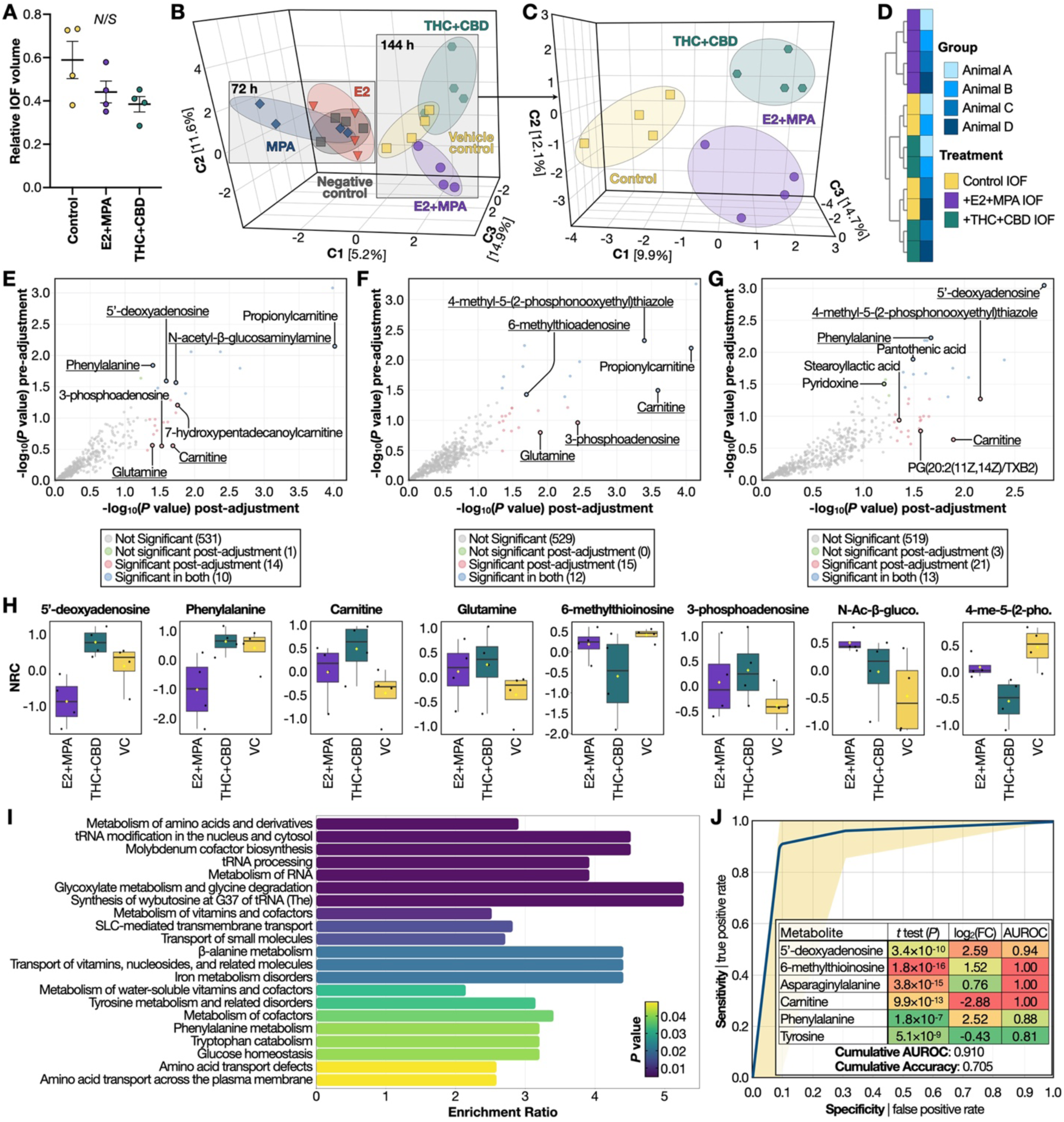
Oviduct organoid secretion composition is disrupted by exocannabinoid exposure. **(A)** Mean (±SD) organoid (n=4) intra-organoid fluid (IOF) relative volumes. No treatment effects observed [not significant (*N*/*S*)]. (**B**) Sparse partial least squares discriminant analysis (sPLS-DA) of organoid IOF metabolomes across controls, 17β-estradiol (E2), medroxyprogesterone acetate (MPA), E2+MPA, and E2+MPA plus Δ⁹-tetrahydrocannabinol (THC) and cannabidiol (CBD) supplementation (THC+CBD). Separation by culture duration also highlighted. (**C**) sPLS-DA of control, E2+MPA, and THC+CBD treated organoids. (**D**) IOF metabolomic profile metadata heatmap with hierarchical clustering by animal and treatment. (**E**-**F**) Linear models with covariate (group) adjustment comparing IOF metabolomic profiles – (**E**) control *vs*. E2+MPA, (**K**) control *vs*. THC+CBD, and (**L**) E2+MPA *vs*. THC+CBD groups. (**M**) Boxplots of select metabolite normalized relative concentrations (NRC) across treatments. Central horizontal lines represent the median value with outer box boundaries depicting upper and lower quartile limits. Error bars depict the minimum and maximum distributions, with yellow rhombi representing the mean values. (**I**) Pathway enrichment analysis using differentially abundant metabolites between E2+MPA *vs*. THC+CBD treated organoid IOF. (**J**) Receiver operating characteristic (ROC) curves generated by input of 6 metabolites (insert) to predict IOF metabolite biomarkers between E2+MPA *vs*. THC+CBD treated organoid IOF. Individual AUROC, t-test *P* values, and fold change (FC) values also provided.

As previously, hierarchical clustering demonstrated a strong treatment effect, with inter-animal variability remaining a secondary contributor (**Fig**. **4D**). Linear models adjusting for this minor batch effect identified 24 DAM between control *vs*. E2+MPA (**Fig**. **4E**), 27 DAM between control *vs*. THC+CBD (**Fig**. **4F**), and 34 DAM between E2+MPA *vs*. THC+CBD groups (**Fig**. **4G**). The most pronounced DAM was 5′-deoxyadenosine, which was elevated in the THC+CBD group (**Fig**. **4H**). Pathway enrichment analysis further revealed intra-organoid fluid DAM associated with amino acid metabolism and RNA processing pathways between E2+MPA *vs*. THC+CBD treated organoids (**Fig**. **4I**).

Statistical significance does not inherently imply predictive utility^61–64^, and although biomarker prediction was not an objective here, we employed a well-established machine learning approach – receiver operating characteristic (**ROC**) analysis – to further evaluate the extent of metabolomic divergence between treatment groups^65^. In this context, the area under the ROC curve (**AUROC**) served as a threshold-independent metric of multivariate separability, reflecting how reliably metabolite profiles distinguished between E2+MPA *vs*. THC+CBD treated organoids. AUROC values can be generally classified as excellent (0.9-1.0), good (0.8-0.9), fair (0.7-0.8), poor (0.6-0.7), or fail (0.5-0.6)^66^.

Interestingly, a minimal panel of just six metabolites achieved high classification performance, accurately distinguishing intra-organoid fluid samples between the two treatment groups with strong sensitivity and specificity (**Fig**. **4J**). Moreover, in this limited metabolite set, predictive accuracy generally aligned with statistical significance, though not necessarily with fold change magnitude. Nevertheless, these findings support the conclusion that exocannabinoid exposure induces distinct, treatment-specific metabolic alterations in organoid secretions.

### Functional impact of altered intra-organoid fluid composition on embryo development

Finally, to determine whether THC+CBD can disrupt embryonic development via direct and/or indirect pathways, we first confirmed CB2 expression in bovine preimplantation embryos (**Fig**. **5A**). CB1 signal absence (not shown) is consistent with bovine embryo transcriptomic studies^21–23^. Embryos were then cultured under four conditions: (*a*) control, (*b*) THC+CBD, (*c*) 5′-deoxyadenosine, or (*d*) THC+CBD+5′-deoxyadenosine (combinatorial), with supplementation during the embryonic genome activation window (**Fig**. **5B**).

**Figure 5.**
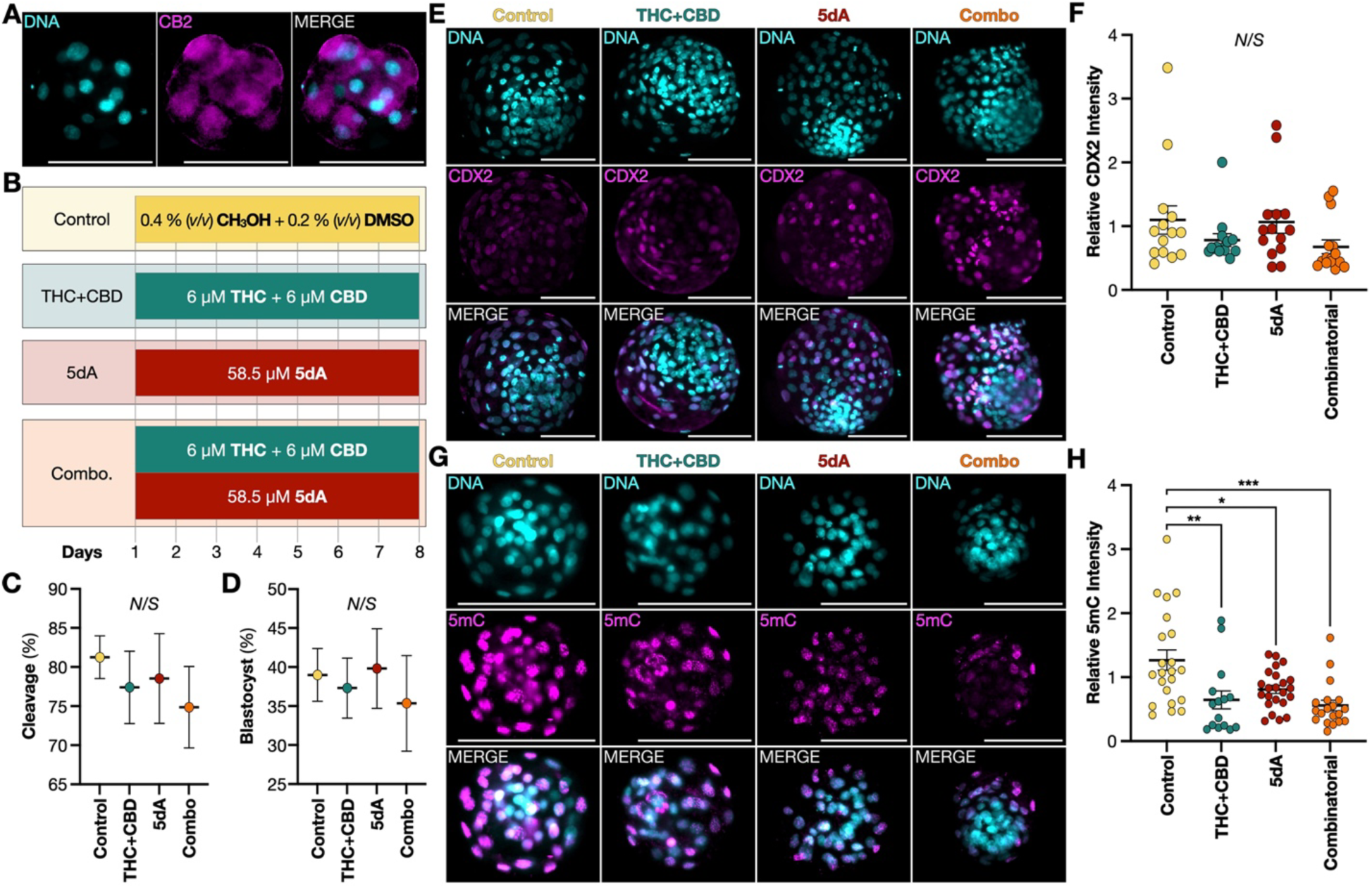
Bovine embryo methylation is disrupted by 5’-deoxyadenosine. (**A**) Representative *in vitro*-derived Day 5 bovine embryo cannabinoid receptor 2 (CB2) expression. (**B**) Independent experimental variables – (*i*) methanol (CH3OH) plus dimethyl sulfoxide (DMSO) control, (*ii*) Δ⁹-tetrahydrocannabinol (THC) plus cannabidiol (CBD), (*iii*) 5’-deoxyadenosine (5dA), and (*iv*) THC+CBD+5dA [combinatorial (combo)] supplementation to embryo culture medium for 192 h. (**C**) Mean (±SEM) embryo cleavage percentage by treatment. No treatment effects observed (n=3) [not significant (*N*/*S*)]. (**D**) Mean (±SEM) embryo blastocyst percentage by treatment (n=3). (**E**) Representative embryo CDX2 expression by treatment. (**F**) Corresponding mean (±SEM) relative CDX2 fluorescence intensity by treatment (n=3). (**F**) Representative 5-methylcytosine (5mC) presence by treatment. (**H**) Corresponding mean (±SEM) relative 5mC fluorescence intensity by treatment (n=3). Select differences observed [*P*≤0.05 (*), *P*≤0.01 (**), and *P*≤0.001 (***)]. All scale bars: 100 µm.

It is worth noting that treatments were administered after fertilization (Day 0) to eliminate potential confounding effects of THC and/or CBD on sperm^67,68^. To additionally control for sexually dimorphic embryo epigenetic responses^69^, sex-sorted (female) sperm was used throughout. Moreover, the 5′-deoxyadenosine concentration [58.5 µM] was selected based on a prior report estimating human extracellular deoxyadenosine levels at approximately 5 µM^70^. This baseline was then scaled by 11.7 – reflecting the observed fold increase in 5′-deoxyadenosine in intra-organoid fluid from THC+CBD *vs*. E2+MPA treated organoids. No significant differences were observed in cleavage (**Fig**. **5C**) or blastocyst formation rates (**Fig**. **5D**), aligning with recent clinical data indicating that cannabis use does not overtly impair fertility^71^. Moreover, blastocyst CDX2 expression, a trophectoderm lineage marker^72^, was unaltered across conditions (**Fig**. **5E**-**F**), suggesting no impact on lineage specification. However, immunofluorescence quantification of 5-methylcytosine revealed reduced DNA methylation in all three treatment groups relative to control (**Fig**. **5G**-**H**), with the most prominent demethylation observed in the combinatorial treatment group.

## DISCUSSION

Oviduct fluid is a dynamic and complex medium composed of ions^73^, metabolites^74^, proteins^75^, and other bioactive factors^76^, primarily shaped by oviduct secretory epithelial cell secretions and vascular transudate exchange^77^. While the functional importance of oviduct fluid in supporting fertilization and early embryo development is well recognized, the physiological mechanisms governing its composition – and their vulnerability to systemic pathological stimuli – remain poorly understood. Furthermore, the developmental consequences of a suboptimal oviduct fluid environment, particularly its potential to influence embryonic trajectory and influence long-term offspring health, are not yet completely defined.

Here we established and characterized bovine oviduct epithelial organoids, which demonstrated appropriate responsiveness to steroid hormones, and whose intra-organoid fluid metabolome closely recapitulated *in vivo* derived oviduct fluid composition. Subsequent exposure to THC+CBD altered organoid morphology, reducing average area without affecting total count, relative to E2+MPA-treated counterparts. This may stem from the observed downregulation of key cell cycle regulators (*MASTL*^78^ and *SASS6*^79^), and/or from versican (*VCAN*) upregulation – an anti-adhesive proteoglycan secreted by epithelial cells^80^ – which may promote organoid detachment from their supporting hydrogel matrix. Additionally, the observed elevated expression of pyruvate dehydrogenase kinase (*PDK2*), which inhibits pyruvate dehydrogenase activity^81^, may indicate a metabolic shift away from oxidative phosphorylation, reducing ATP availability for proliferation – a highly energy-demanding process^82^.

Subsequent transcriptomic and secretomic profiling revealed that THC+CBD exposure altered the organoid secretome, including elevated intra-organoid fluid concentrations of 5′-deoxyadenosine. More specifically, THC+CBD appeared to dampen the type I interferon-mediated^83^ stress response^84^, as evidenced by downregulation of *HERC5*^55^, *IDO1*^56^, *CXCL10*^57^, *RIGI*^58^, *STOML1*^59^, and *RSAD2* (viperin)^60^. *RSAD2* dysregulation in the endometrium has previously been associated with impaired uterine receptivity in ovine^85^ and murine^86^ models.

While viperin and related radical S-adenosylmethionine (**SAM**) enzymes catalyze SAM to 5′-deoxyadenosine as a byproduct^87,88^, the paradoxical increase in 5′-deoxyadenosine, despite *RSAD2* downregulation, may instead reflect upregulation of xenobiotic metabolism pathways, including *UGT1A6*, *CYP1A1*, and *CYP1B1*. The latter can generate deoxyadenosine adducts^89^, likely underpinning the 5′-deoxyadenosine accumulation observed here. It is also tempting to suggest that THC+CBD treatment shifted 5′-deoxyadenosine export from vesicular mediated secretion (*e*.*g*., *ATP8B4* downregulation)^90^ toward transporter-mediated efflux, given the increased expression of several *SLC25* family mitochondrial carriers. Although further validation, including at the protein level, is needed, this may represent an adaptive mechanism to maintain cellular homeostasis under xenobiotic stress.

As expected, direct exocannabinoid exposure induced embryonic DNA hypomethylation^91,92^. More novel, however, is our observation that exposure to 5′-deoxyadenosine, an endogenous metabolite, also appears to reduce embryonic DNA methylation. Although studies on 5′-deoxyadenosine in mammalian systems are limited, it is known to be taken up by mammalian cells^93^ and can inhibit radical SAM enzyme activity^94^. Therefore, the mechanism by which 5′-deoxyadenosine disrupts embryonic methylation likely involves competitive inhibition of methionine adenosyltransferase (**MAT**), reducing conversion of methionine to SAM and thereby lowering available substrate for methyltransferase activity^95^. In other words, MAT inhibition by 5′-deoxyadenosine would reduce the SAM:SAH ratio, leading to global hypomethylation – a phenomenon we observed experimentally. This may explain, at least in part, the link between maternal prenatal cannabis and DOHaD-associated phenotype development^12^.

Future research directions include validating these findings in human oviductal epithelial organoids, potentially using stromal-epithelial co-cultures to enhance physiological relevance^96–98^, and testing developmental outcomes using human blastoids^99^. Determining whether 5′-deoxyadenosine accumulates *in vivo* following maternal exocannabinoid exposure, especially in murine models, would further clarify physiological relevance. Moreover, mapping genome-wide methylation changes following 5′-deoxyadenosine exposure would identify the gene regulatory networks disrupted by 5′-deoxyadenosine. Beyond intra-organoid fluid metabolomics, proteomic and extracellular vesicle profiling^100^ would shed further light on maternal-embryo crosstalk mechanisms during this sensitive developmental window.

Furthermore, GPR55 (a putative cannabinoid receptor)^101^ and TRPV1 (a CBD-responsive cation channel)^102^ are expressed in epithelial tissues. Investigating their potential roles in mediating exocannabinoid effects on the oviduct may offer a more comprehensive understanding of the molecular mechanisms underlying our findings. Finally, it is important to note that our metabolomic screen did not detect THC or CBD within intra-organoid fluid, suggesting that they do not transverse the oviduct epithelial monolayer. Although *in vitro* and *in vivo* targeted metabolomics approaches are required to confirm this, our data strongly suggest that exocannabinoids THC and CBD exert indirect effects on the embryo by modulating maternal oviduct physiology rather than via direct embryo exposure.

In summary, we established bovine oviduct epithelial organoids as a robust model for sequentially examining how maternal exposures shape luminal fluid composition and downstream developmental outcomes. We demonstrate that exocannabinoid exposure alters epithelial morphology, transcriptomic profiles, and secretory function, notably increasing 5′-deoxyadenosine secretion – a metabolite capable of inducing embryonic DNA hypomethylation. These findings underscore the sensitivity of early developmental environments to systemic xenobiotic exposure and highlight the value of oviductal organoids in elucidating mechanisms underlying developmental programming. More broadly, these data highlight the utility of organoids for interrogating the dynamic regulation of luminal fluid composition, which could extend to cerebrospinal, synovial, intraocular, follicular, uterine, and other interstitial fluids.

While exocannabinoids were used here as a model, this system can be broadly applied to other compounds of concern within the DOHaD framework.

## MATERIALS AND METHODS

### Sample generation

All animal-related procedures were approved by the Louisiana State University Agricultural Center Institutional Animal Care and Use Committee (IACUC) protocols A2022-11 and A2022-24. The estrous cycles of four crossbred cattle, with a mean (± SD) age of 8.5 ± 2.1 years and weight of 445 ± 58.7 kg, were synchronized using a standard protocol^103,104^, involving intramuscular administration of a gonadotropin-releasing hormone (**GnRH**) analogue (Fertagyl; Merck Animal Health, 014148) alongside intravaginal insertion of a P4 (1.38 g) controlled internal drug release (**CIDR**) device (Eazi-breed; Zoetis Animal Health, 021274). Seven days later, a prostaglandin F2α (**PGF2α**) analogue (Lutalyse; Zoetis Animal Health, 026256) was administered intramuscularly, followed by CIDR removal the following day. Estrous detection was performed using breeding indicators (Estrotect; Rockway, 78013). Indicators were monitored twice daily starting 48 h following CIDR withdrawal. All animals displayed signs of standing estrous and were sacrificed on Day 5 of metestrus (**Fig**. **1A**).

### Sample collection

Complete reproductive tracts from synchronized cattle (n=4) were obtained at a local commercial abattoir (Coastal Plains Meat Company, Eunice, LA), immediately placed in sealed plastic bags on ice, and transported to the laboratory within 1.5 h. Upon arrival, the presence of a stage-appropriate ovarian *corpus luteum* was confirmed, and the corresponding ipsilateral oviduct was carefully excised. One quarter of each oviduct isthmic region was then separated, placed in aqueous 4 % (*v*/*v*) formaldehyde (VWR, 10790-712) in Milli-Q [18.2 MΩ·cm] water and kept at 4 °C until downstream immunohistochemical analysis.

Oviduct fluid was then flushed by careful insertion and clamping of a 20 G needle into the oviduct isthmus, followed by gentle manual expulsion of 2 ml 1x phosphate buffered saline (**PBS**) (Fisher Bioreagents, BP39920) throughout the remaining oviduct. Oviduct fluid was aliquoted, snap frozen in liquid nitrogen [**N_2_(*l*)**], stored at -80 °C overnight, and subsequently stored in N_2_(*l*) until analysis. Following oviduct fluid collection, each oviduct was manually squeezed from isthmus to infundibulum to expel the cellular homogenate into 100 mm culture dishes (Thermo Scientific, 263991) containing Dulbecco’s Modified Eagle Medium (**DMEM**)-F12 medium (Gibco-Invitrogen, 11320033), supplemented with 10 % (*v*/*v*) antibiotic-antimycotic solution (Gibco-Invitrogen, 15240062), similarly to previously^105^. This cell suspension was then transferred into conical tubes (Falcon, 352196) and centrifuged at 300 × *g* for 10 min at 4 °C. Following centrifugation, the cell pellet was divided into two equal portions – one was preserved in RNA-later (Invitrogen, AM7021) and stored at -80 °C until processing for bulk RNA-seq, whereas the other was immediately processed for organoid establishment.

### Tissue immunohistochemical labelling

As aforementioned, one quarter of each oviduct isthmus was submerged in aqueous 4 % (*v*/*v*) aqueous formaldehyde and kept at 4 °C for no longer than one week. Tissues were then washed three times in PBS and transferred into 30 % (*v*/*v*) sucrose (Fisher Bioreagents, BP220-212) in PBS at 4 °C for approximately 1 month. Thereafter, tissues were embedded in Tissue-Tek^®^ Optimal Cutting Temperature compound (Finetek, 4583) within Tissue-Tek^®^ plastic cryomolds (Finetek, 4566) and stored at -80 °C for at least 6 h. Embedded tissue blocks were cryo-sectioned (Thermo Scientific, HM525NX) at a thickness of 5 µm, mounted onto charged slides (VWR, 48311-703), and stored at -20 °C. For immunohistochemistry, slides were first incubated (37 °C) for 5 min before further fixation by submersion in 4% (*v*/*v*) formaldehyde in Milli-Q water for 10 min at room temperature (**RT**) with orbital shaking. Slides were then washed three times in PBS (10 min each) and twice in Milli-Q [18.2 MΩ·cm] water (2 min each).

Antigen retrieval was achieved by incubation in pre-heated 10 % (*v*/*v*) aqueous Reveal Decloaker solution (Biocare Medical, RV1000MMRTU) at 98 °C for 30 min. Slides were cooled by incubation at RT for 1 h before washing twice in PBS (5 min each) with orbital shaking at RT. For blocking, a hydrophobic barrier was drawn around the tissue (Electron Microscopy Sciences, 71310) and 10 % (*v*/*v*) Normal Goat Serum (**NGS**) (AbCam, ab7481) in PBS was applied. Slides were incubated for 1 h at RT in a humidified, dark chamber (Electron Microscopy Sciences, 71397-B). Primary antibodies (**Table S1**) were diluted to 1 µg·ml^-1^ in 10 % (*v*/*v*) NGS in PBS and applied to slides overnight at 4 °C in a humidified, dark chamber. Slides were then washed three times in PBS (10 min each). The secondary antibody (**Table S1**) was also diluted to 1 µg·ml^-1^ in 10 % (*v*/*v*) NGS in PBS, and applied to slides for 1 h at RT, followed by three PBS washes (10 min each). Nuclear counterstaining was performed using Hoechst 34580 (AAT Bioquest, 17537), diluted 1:5000 in PBS, and applied to slides for 5 min at RT, followed by three PBS washes (5 min each). Finally, sections were mounted with Fluoromount-G (Invitrogen, 00-4958-02) and covered (VWR, 48404-453). Imaging was performed using a Leica DMI8 fluorescence microscope coupled to Leica Application Suite X (LASX) software (v. 3.9.128433). Images are provided in **Figure 1C**.

### Bovine oviduct epithelial organoid establishment and culture

Each cell suspension was resuspended in chilled Wash Medium (**Table S2**) before centrifugation (300 × *g* for 10 min at 4 °C) and resuspension in pre-equilibrated (37 °C) Stromal Medium (**Table S2**) and transfer to one 75 cm^2^ flask (VWR, 10062-860) per oviduct. Cell homogenates were cultured at 38.5 °C under 5 % CO_2_ in humidified air for 18 h to achieve differential attachment of stromal and epithelial cells^106^. Thereafter, the conditioned medium from each oviduct was transferred to a conical tube and centrifuged as above to pellet the epithelial cell population. The pellet was resuspended in 1 ml chilled Wash Medium, counted using trypan blue (VWR, K940) exclusion using an automated cell counter (Corning, 6749), before resuspension in undiluted reduced growth basement membrane (Cultrex) hydrogel (R&D Systems, 3433005R1) at a volume achieving a dilution of 500 cells·µl^-1^. Thereafter, working on ice, cells were transferred to 12 well plates (Fisherbrand, FB012928) at a density of four 20 µl hydrogel domes per well, before incubation at 38.5 °C under 5 % CO_2_ in humidified air for 30 min to facilitate hydrogel polymerization. Each well was then flooded with 600 µl pre-equilibrated (37 °C) Thawing Medium (**Table S2**).

Cultures were maintained at 38.5 °C under 5 % CO_2_ in humidified air with medium replenishment every 48 h for 5 days, by which point organoids began to form. More specifically, Thawing Medium was used for the first two media changes, after which Expansion Medium (**Table S2**) was used for all subsequent media changes. To evaluate capacity for cryopreservation, organoids were first removed by replacing conditioned Expansion Medium with chilled Wash Medium before manually dislodging the hydrogel using a wide-orifice 1000 µl pipette tip (Thermo Scientific, 02707008). The cell-hydrogel suspension was transferred to a conical tube, centrifuged as above, and resuspended in chilled Wash Medium. This wash step was repeated until the hydrogel was no longer visible by eye.

The resulting pellet was resuspended in Freezing Medium (**Table S2**) and stored at -80 °C within a freezing container (Mr. Frosty, Thermo Scientific, 5100-0036), to achieve a cooling rate of approximately 1 °C·min^-1^. The following morning, cells were stored in N_2_(*l*). Following 3 months, organoids were thawed by gradual addition of pre-equilibrated (37 °C) Thawing Medium (**Table S2**) and gentle mixing. Suspensions were then transferred to conical tubes and centrifuged as above. The supernatant was discarded, and the pellet resuspended thoroughly in chilled Wash Medium. Thereafter, cell counting, centrifugation, Cultrex hydrogel resuspension, plating, and incubation in Expansion Medium, was performed as described above.

### Oviduct organoid hormonal and exocannabinoid supplementation

Following 14 days of culture, organoids from each animal were assigned to one of six groups, as schematically depicted in **Figures 1C** and **3C**. These were: (*a*) negative control – comprising Base Medium (**Table S2**); (*b*) E2 – Base Medium supplemented with E2 (Thermo Scientific Chemicals, L0380103) dissolved in methanol (Fisher Chemical, A4521) at a final concentration of 50 nM with a corresponding vehicle contribution of 0.6 % (*v*/*v*); (*c*) MPA – Base Medium supplemented with MPA (Thermo Scientific Chemicals, 461120010) dissolved in methanol at a final concentration of 1 µM with a corresponding vehicle contribution of 0.3 % (*v*/*v*); (*d*) Physiological mimic – Base Medium supplemented with both E2 and MPA as above, with a corresponding cumulative vehicle contribution of 0.9 % (*v*/*v*); (*e*) Pathological mimic – Base Medium supplemented with E2 and MPA as above, in addition to THC (Cayman Chemical, ISO60157) dissolved in methanol at a final concentration of 6 µM with a corresponding vehicle contribution of 0.2 % (*v*/*v*); and CBD (Cayman Chemical, ISO60156) dissolved in methanol at a final concentration of 6 µM with a corresponding vehicle contribution of 0.2 % (*v*/*v*); and (*f*) vehicle control – Base Medium supplemented with 1.3 % (*v*/*v*) methanol – identical to the largest solvent contribution (Group *e*). Media was replenished every 24 h throughout the treatment period. Groups *a*-*c* and *d*-*e* were cultured for 3 and 6 days, respectively.

### Oviduct organoid morphological assessment

Organoids imaged by brightfield microscopy at 5x magnification (Leica DMI8) every 48 h during the 14-day expansion period, and every 24 h during treatment. Thereafter a single dome from each animal and treatment group was selected at random for analysis using Fiji (ImageJ, National Institutes of Health) with the SAMJ plugin. Specifically, the *points* function within the *Efficient SAM* module was used to manually select all organoids in each image to quantify the count, circularity, and area of each individual organoid. Representative images are provided in **Figures 1I** and **3D**. Means and standard errors were calculated using Microsoft Excel for Mac (v. 16.97.2, b. 25052611) and plotted using GraphPad Prism for Mac version (v. 10.4.2, b. 534) (**Fig**. **1F-H** & **3E-G**). Statistical details are described below.

### Intra-organoid fluid extraction

We utilized the previously established high-throughput centrifugation protocol for extracting intra-organoid fluid^37^. Briefly, at the time of organoid harvest, the conditioned Base Medium in each well was replaced with chilled Wash Medium before manually dislodging the hydrogel using a wide-orifice pipette tip. The cell-hydrogel suspension was transferred to a conical tube, centrifuged (300 × *g* for 10 min at 4 °C), and resuspended in chilled Wash Medium. This wash step was repeated until the hydrogel was no longer visible by eye. At this point, the pellet was resuspended in chilled PBS and re-centrifuged as above. This wash step was repeated twice, by which point the PBS was completely clear, and organoids were resuspended in 100 µl chilled PBS. Organoids were then transferred to a 1.5 ml tube (VWR, 89000-028) and centrifuged at 4,000 × *g* for 30 min at 4 °C. The volume of diluted intra-organoid fluid supernatant was recorded, and intra-organoid fluid was transferred to a 1.5 ml tube, snap frozen [N2(*l*)] and stored at -80 °C overnight, before transfer to N_2_(*l*) until analysis. Intra-organoid fluid volumes (**Fig**. **2F** & **4A**) are presented as a ratio of the volume recovered to the total RNA concentration extracted from the residual organoid pellet, described below, to account for variation in organoid mass.

### Oviduct RNA extraction

Immediately following intra-organoid fluid isolation, RNA was extracted from each residual organoid pellet using the PureLink™ RNA Mini Kit (Invitrogen, 12183018A) in accordance with manufacturer instructions. In brief, organoids were first lysed by addition of Lysis Buffer supplemented with 2-mercaptoethanol (Millipore-Sigma, 444203250ML). The lysate was thoroughly mixed by vortexing. An equal volume of aqueous 70 % (*v*/*v*) ethanol (Koptec, V1001) was then added to the lysate. The mixture was loaded onto a spin cartridge and centrifuged at 12,000 × *g* for 15 sec at RT. The flow-through was discarded, and the cartridge washed with Wash Buffer I once and Wash Buffer II twice, with centrifugation as above following each wash. After the third wash, the spin cartridge was further centrifuged (12,000 × *g* for 2 min at RT) to remove residual Wash Buffer. RNA was eluted by addition of 30 μl DEPC-treated RNase free water (Thermo Scientific, R0601) to each spin cartridge before incubation at RT for 1 min. Each column was then centrifuged (12,000 × *g* for 2 min at RT) for RNA collection. RNA yield and purity were determined using a NanoDrop One spectrophotometer (Thermo Fisher Scientific). RNA was stored at -80 °C until analysis. RNA concentrations recovered from each sample are provided in **Table S3**.

### cDNA library preparation

A total of 200 ng RNA per sample was processed using a single cell/low input RNA library preparation kit (New England Biolabs, E6420L), in accordance with manufacturer instructions. In brief, reverse transcription was performed using a template-switching mechanism to synthesize full-length cDNA, followed by amplification with 8 polymerase chain reaction (**PCR**) cycles to yield sufficient material. Thereafter, amplified cDNA was purified using ‘DINOMAG’ solid-phase reversible immobilization (**SPRI**) Select beads (Redoxica, DN9004), before fragmentation and end-repair using a fragmentation system (**FS**) kit (New England Biolabs, E7805L) in line with manufacturer instructions. In brief, adapters were ligated, and cDNA libraries were enriched via 8 PCR cycles using multiplex oligos for dual index primers (New England Biolabs, E6446S). Final libraries were quantified using a Qubit 4 (Thermo Fisher Scientific, Q33238) fluorometric assay and assessed for size distribution and quality on an Agilent Bioanalyzer (Agilent Technologies), using corresponding sample buffers (Agilent Technologies, 5067-5589) and screen tape (Agilent Technologies, 5067-5588). Prepared libraries were stored at -20 °C until sequencing.

### RNA sequencing

RNA-seq was outsourced to Novogene, where libraries were sequenced using the Illumina NovaSeq X Plus platform with paired-end 150 base pair reads (PE150).

### Transcriptomic analysis

Raw transcript read counts (**Table S4**) were processed primarily using the integrated differential expression and pathway analysis (**IDEP**) 2.01 platform. More specifically, raw read counts were uploaded and aligned to the *Bos taurus* genome ENSEMBL assembly (ARS-UCD1.2). IDEP confirmed the presence of 32,607 transcripts across 24 samples, of which 27,482 passed initial filtration. Among these, 17,839 were assigned an ENSEMBL gene ID, while the remaining 9,643 retained their original ID and were included in downstream analysis. Subsequent processing steps included the exclusion of one sample (MPA-D) to avoid sequencing depth bias, due to insufficient sequencing depth (< 150,000 raw reads). Thereafter, data were uploaded with individual groups for each analysis using imputation of missing values using the group median. PCA (**Fig**. **2A**) was performed using the IDEP *PCA 3D* function.

Heatmaps (**Fig**. **2B** & **3I**) were generated using the k-means clustering function within IDEP, applying the default setting of six clusters and selecting the top 10,000 DEG, with gene-wise centering (mean subtraction) applied. For each cluster, corresponding Gene Ontology Biological Process enrichment data were downloaded and used to construct multivariate bubble plots (**Fig**. **2C** & **3J**) in GraphPad Prism. Bubble size in these plots represents the impact ratio, calculated as the number of DEG associated with a given pathway divided by the total number of genes annotated to that pathway. Individual DEG volcano plots were generated in IDEP using the DEG2 function, employing the DESeq2 method with a false discovery rate (**FDR**) threshold of 0.1 and a minimum fold change of 2.0. Independent filtering of low-count genes was applied. Resulting data were exported and re-plotted (**Fig**. **2D** & **3K-M**) in GraphPad Prism. Parametric gene set enrichment analysis was also performed within IDEP using the Pathway function, with an FDR cutoff of 0.1 and referencing the GO Biological Process and/or GO Molecular Function databases (**Fig**. **2E**&**3N**).

### Mass spectrometry

High-throughput, untargeted metabolomic profiling of intra-organoid fluids was performed at the Louisiana State University (**LSU**) Mass Spectrometry Facility. Prior to analysis, intra-organoid fluid was thawed on ice for 60 min. Thereafter 60 µl of each sample was supplemented with 120 µl methanol and 120 µl acetonitrile. Tubes were vortexed for 15 sec, sonicated for 5 min, and incubated at -20 °C for 1 h, followed by centrifugation (15,000 × *g* for 10 min at 4 °C).

Resulting supernatants were analyzed using a Synapt XS Electrospray Ionization Quadrupole– Ion Mobility Separation–Time-of-Flight (ESI-Q-IMS-TOF) mass spectrometer (Waters Corporation, RRID: SCR-026524) coupled to an Acquity Premier Ultra-Performance Liquid Chromatography (**UPLC**) system (Waters Corporation, RRID: SCR-027039). Samples were analyzed in both positive (capillary voltage: 1.5 kV) and negative (2.0 kV) electrospray ionization modes across a mass range of 50–1200 m/z. Lockmass calibration was achieved using a 200 pM leucine-enkephalin (Leu-Enk) solution (Waters, 186006013), introduced every 10 sec via a secondary orthogonal sprayer. The system operated in data-independent acquisition with elevated energy (**MSE**) mode with a 0.25 sec acquisition time per scan and a collision energy ramp of 10–30 V. Chromatographic separation was performed using a Premier Bridged Ethylene Hybrid (**BEH**) Amide column (100 × 2.7 mm, 1.7 µm pore size) (Waters, 186009508) fitted with a matching Vanguard guard column (5 mm length) (Waters, 186009510).

The mobile phase consisted of Buffer A [0.1 % (*v*/*v*) aqueous formic acid (Fisher Scientific, A117-50)] and Buffer B [0.1% (*v*/*v*) formic acid in acetonitrile (Millipore Sigma, 900682)] at a constant flow rate of 400 µl·min^-1^. The gradient program was as follows: 5% Buffer A (0–1 min), 95% Buffer A (1–9 min), and 5% Buffer A (9–11 min). A pooled quality control (**QC**) sample was prepared prior to each batch to optimize sample dilution, monitor instrument performance, and serve as a reference for peak alignment. Injection volumes were 1 µl in positive mode and 2 µl in negative mode.

Data acquisition and initial review were conducted using MassLynx version 4.2 (Waters Corporation). Files were imported into Progenesis QI (v. 3.0) for alignment, peak detection, and mass deconvolution, using standard parameters. Metabolite identification was first performed using the Metabolite and Tandem MS Database (**METLIN**) MS/MS library (2019). Remaining unidentified features were matched against the Human Metabolome Database (**HMDB**) using in silico fragmentation prediction^107^.

### Metabolomic analyses

A total of 1,717 metabolite features were identified. All raw peak areas (**Table S5**) were then normalized to the total RNA recovered in organoid pellets following intra-organoid fluid isolation (**Table S3**) to account for organoid mass. Single-factor analyses were conducted using MetaboAnalyst 6.0^108^. Initially, 65 metabolites with a constant or single value across samples were removed as part of data integrity checking. The raw data were then normalized to the median, log-transformed (base 10), and pareto-scaled (mean-centered and divided by the square root of the standard deviation of each variable).

Principal component analysis (**PCA**) plot (**Fig**. **2G**) with 95% confidence intervals was created using permutational multivariate analysis of variance (**PERMANOVA**), with distributions based on Euclidean distance from the first two principal components. Sparse partial least squares discriminant analysis (**sPLS**-**DA**) was performed with 5 components and 10 variables per component. Model performance was evaluated using 5-fold cross-validation with an increasing number of components and a fixed 10 variables per component (**Fig**. **2H** & **4B-C**).

For metadata (*i*.*e*., batch correction) analyses, peak intensities were filtered, normalized, transformed, and scaled as described above. Metadata heatmaps with hierarchical clustering (**Fig**. **2I** & **4D**) were generated using Euclidean distance and Ward clustering for both metabolites and metadata variables. Linear models with covariate adjustment (*P*≤0.05) were performed within MetaboAnalyst 6.0, using the *limma* linear regression approach, as previously described (**Fig**. **2J-L** & **4E-G**)^109,110^. Corresponding boxplots of specific metabolite normalized relative concentrations (**Fig**. **2M** & **4H**) were exported from MetaboAnalyst.

Categorical (classification) pathway impact analysis (**Fig**. **2N**) was conducted by first standardizing compound names against the HMDB, PubChem, and KEGG databases, before peak intensity normalization, transformation, and scaling – all as above. The pathway analysis focused on significant metabolites rather than pre-selected ones, with enrichment performed using the *global test* as above. Topology was assessed using relative-betweenness centrality, and the reference metabolome included all compounds from the KEGG *Bos taurus* library. A scatter plot was generated to display all matched pathways, with *P*-values from the pathway enrichment analysis plotted against pathway impact values from the topology analysis.

For categorical (classification) quantitative enrichment analyses (**Fig**. **4I**), compound names were first standardized against the HMDB, PubChem, and KEGG databases. Unstandardized compound names were excluded. Peak intensities were then normalized, transformed, and scaled as in prior steps. Enrichment testing was conducted based on the *global test*^111^ against the RaMP-DB metabolite set library, which integrates 3,694 features from KEGG (via HMDB), Reactome, and WikiPathways databases. Only metabolite sets with at least two entries were included. The enrichment ratio was calculated by comparing the number of observed metabolites to the expected number within each set.

### Machine-learning based biomarker identification

Biomarker analysis (**Fig**. **4J**) was also performed using MetaboAnalyst 6.0, leveraging the ROC curve-based model evaluation function. Initially, raw peak intensities were filtered, normalized, transformed, and scaled as described above. Metabolites were then manually selected for ROC analysis, which was conducted using the *Random Forests* multivariate algorithm. Specifically, 100 cross-validations were performed, with results averaged to generate a ROC curve, with 95% confidence intervals and a predictive accuracy values also presented.

### Oviduct organoid whole mount immunohistochemical labelling

Following treatment, organoids were isolated by replacing the culture medium from each well with 600 µl VitroGel Cell Recovery Solution (The Well Bioscience, MS03-100), before incubation at 4 °C for 30 min to dissolve the hydrogel. Organoids were then transferred to conical tubes and centrifuged (300 × *g* for 4 min at 4 °C). The supernatant was discarded, and the pellet resuspended in 2 ml Wash Medium. This wash step was repeated until the hydrogel was no longer visible by eye. Organoids were then resuspended in 4 % (*v*/*v*) aqueous formaldehyde for 20 min at RT with orbital shaking. Thereafter, organoids were centrifuged as above, resuspended in 1 ml PBS and kept at 4 °C for no longer than one week. Organoids were then centrifuged as above and permeabilized by incubation with 0.1 % (*v*/*v*) Triton X-100 (Apacor, 1499) in PBS for 30 min at RT with orbital shaking. Following centrifugation as above, blocking was achieved by incubation with 5 % (*w*/*v*) bovine serum albumin (**BSA**) (Sigma Aldrich, A6003) in PBS for 1 h at RT with orbital shaking.

Organoids were centrifuged as above, then incubated overnight at 4 °C with the primary antibody (**Table S1**) diluted to 1 µg·ml^-1^ in 10 % (*v*/*v*) NGS in PBS. They were washed three times – each wash involving centrifugation and resuspension in 0.1 % (*v*/*v*) BSA and 0.1 % (*v*/*v*) Tween-20 (Sigma-Aldrich, P7949) in PBS – before incubation for 1 h at RT with the secondary antibody (**Table S1**) also diluted to 1 µg·ml^-1^ in 10 % (*v*/*v*) NGS in PBS. After two further PBS washes and centrifugation as above, organoids were resuspended in Hoechst (1:5000 in PBS) for 10 min at RT with orbital shaking, washed three more times with PBS, and transferred onto slides using an EZ-Grip (Cooper Surgical, 7722802). Finally, organoids were mounted in Fluoromount-G, covered, and imaged as described above.

### In vitro embryo production

Abattoir derived bovine cumulus-oocyte complexes (**COC**) were purchased from Simplot (Meridian, ID, USA) and transported overnight following collection, in a proprietary M199-based maturation medium, within an incubator maintaining 38.5 °C. Twenty-two (22) h following initial COC harvest, *in vitro* fertilization (**IVF**) was performed using a standard protocol^112^. In brief, oocytes were washed four times by manual transfer between wells of a 4-well plate (SPL, 30004) containing BO-IVF Medium (IVF Bioscience, 71004), using a WireTrol II manipulator (Drummond Scientific, 5-000-2050) to remove maturation medium and debris.

Subsequently, groups of approximately 50 COC were transferred into individual wells of 4-well plates, each containing 400 μl BO-IVF medium, and incubated at 38.5 °C under 5 % CO_2_ in humidified air for approximately 5 min. In this time, sperm were prepared. Specifically, female sex-sorted bovine semen straws (ST Genetics, Pristine) were thawed by submersion in 37 °C water for 45 sec, before transferring straw contents to a conical tube containing pre-equilibrated (37 °C) BO-SemenPrep Medium (IVF Bioscience, 71003). Sperm were then centrifuged (300 × *g* for 5 min at RT) and the resulting pellet was resuspended in BO-SemenPrep Medium before centrifugation as previously.

This final pellet was resuspended in pre-equilibrated (38.5 °C under 5 % CO_2_ in humidified air overnight) BO-IVF medium. Sperm concentrations were determined using a hemocytometer to achieve the addition of 2 × 10^6^ spermatozoa per 50 COC. Gametes were co-cultured (38.5 °C under 5 % CO_2_ in humidified air) for 18 h. Following fertilization, presumptive zygotes were denuded of cumulus cells by manual transfer into a microcentrifuge tube containing pre-equilibrated (37 °C) BO-Wash Medium (IVF Bioscience, 51002), followed by gentle vortex for 5 min. Presumptive zygotes were then washed four times by manual transfer between wells of a 4-well plate containing BO-Wash Medium, using a WireTrol II manipulator (Drummond Scientific, 5-000-2005). Washed presumptive zygotes were similarly transferred, in groups of 50, into 4-well plates containing pre-equilibrated (37 °C) BO-IVC Medium (with treatments, as described below), per well. Culture proceeded for 24 h (38.5 °C under 5 % CO_2_ and 5 % O2 under N_2_).

### Embryo exocannabinoid and 5′-deoxyadenosine supplementations

On Day 1 post-fertilization, embryos were manually transferred at random into wells of a 4-well plate corresponding to one of four groups, as depicted in **Fig**. **5B**. These were: (*a*) vehicle control – comprising BO-IVC Medium with 0.4 % (*v*/*v*) methanol and 0.2 % (*v*/*v*) dimethyl sulfoxide (**DMSO**) (VWR, 0231); (*b*) THC+CBD – comprising BO-IVC with THC dissolved in methanol, and supplemented at a final concentration of 6 µM with a corresponding vehicle contribution of 0.2 % (*v*/*v*); and CBD dissolved in methanol, and supplemented at a final concentration of 6 µM with a corresponding vehicle contribution of 0.2 % (*v*/*v*); (*c*) BO-IVC with 5′-deoxyadenosine (Cayman Chemical, 29619) dissolved in DMSO, and supplemented at a final concentration of 58.5 μM with a corresponding vehicle contribution of 0.2 % (*v*/*v*); and (*d*) combinatorial – comprising BO-IVC with 6 µM THC, 6 µM CBD, and 58.5 μM 5′-deoxyadenosine as above, with corresponding vehicle contributions identical to that of the vehicle control.

Treatment plates were prepared and equilibrated (38.5 °C under 5 % CO_2_ and 5 % O2 under N_2_) for 18 h prior to embryo exposure. Treatments were prepared to reach a final volume of 500 μl per well. Following embryo placement, each well was overlaid with light mineral oil (Fujifilm, Irvine Scientific, 9305). Culture proceeded for 8 days (38.5 °C under 5 % CO_2_ and 5 % O2 under N_2_). Cleavage and blastocyst formation were assessed on Days 3 and 8 post-fertilization, respectively. A total of 737 presumptive zygotes were distributed across the four treatment groups and cultured over three independent replicates.

### Embryo whole mount immunohistochemical labelling

Our embryo whole-mount imaging protocol was adapted from Dr. Peter J. Hansen’s laboratory at the University of Florida^113^. Briefly, blastocyst-stage embryos were washed three times in PBS by manual transfer between wells of a 4-well plate containing PBS, using a WireTrol II manipulator. Embryos were then fixed in 4 % (*v*/*v*) paraformaldehyde (**PFA**) (Electron Microscopy Sciences, 15714) diluted in PBS for 20 min at RT, followed by three additional PBS washes as above. Permeabilization was carried out by transferring embryos into 0.5 % (*v*/*v*) Triton X-100 in PBS for 30 min at RT, after which embryos were moved and incubated inro 5 % (*w*/*v*) BSA in PBS for 1 h at RT. Embryos were then transferred into a solution comprising 1 µg·ml^-1^ CDX2 primary antibody (**Table S1**) in 1 % (*w*/*v*) BSA with 0.1% (*v*/*v*) Tween-20 in PBS, and incubated overnight at 4 °C.

Following primary antibody incubation, embryos were washed three times for 2 min each by transfer between wells containing 0.1 % (*w*/*v*) BSA with 0.1 % (*v*/*v*) Tween-20 in PBS. Embryos were then similarly incubated with the secondary antibody (**Table S1**), diluted to 1 µg·ml^-1^ in 0.1 % (*v*/*v*) Tween-20 and 1 % (*w*/*v*) BSA in PBS for 1 h at RT. This was followed by three additional washes using 0.1 % (*v*/*v*) Tween-20 with 0.1 % (*w*/*v*) BSA in PBS for 2 min each. For nuclear counterstaining, embryos were transferred into Hoechst (1:5000 dilution in PBS) for 5 min at RT, then washed three times in Embryo Wash Solution for 2 min each. Finally, embryos were transferred onto microscope slides and mounted using Fluoromount-G. Imaging was performed as described above.

### Embryo whole mount DNA methylation labelling

Immunolabeling of bovine embryo DNA for 5’-methylcytosine was performed following a modified protocol^114^. Briefly, Day 8 blastocysts were collected and washed thoroughly in PBS three times, similarly to previously. Embryos were then transferred into 4 % (*v*/*v*) PFA diluted in PBS for 30 min at RT, followed by three washes in PBS. Permeabilization was carried out by transferrin embryos into 0.5 % (*v*/*v*) Tween-20 with 0.5 % (*v*/*v*) Triton X-100 in PBS at RT for 40 min. Antigen retrieval was achieved by embryo transfer into 4M HCl with 0.1% (*v*/*v*) Triton X-100 at RT for 10 min, followed by three washes in Phosphate Buffered Tween (**PBT**) – 0.05% (*v*/*v*) Tween-20 in PBS. Embryos were then transferred into pre-equilibrated (37 °C for 2 h under mineral oil) 50 µl drops of 0.25% (*v*/*v*) Trypsin-EDTA (Sigma Aldrich, SM-2003-C) for 40 sec at 37 °C. Digestion was terminated by transferring embryos into 10 % (*v*/*v*) NGS in PBT for 2 min at 37 °C, followed by three washes in PBT supplemented with 2 % (*v*/*v*) BSA (**PBSA**).

Embryos were then transferred to 30 % (*v*/*v*) NGS in PBT at RT for 1 h before incubation with 10 μg·ml^-1^ anti-5meC primary antibody (**Table S1**) in 5 % (*v*/*v*) NGS in PBT for 1 h at RT. After three PBSA washes as previously, embryos were incubated for 1 h at RT with the secondary antibody (**Table S1**) diluted 1:300 in PBSA, followed by a further three PBSA washes. Nuclei were counterstained by incubation with Hoechst (1:5000 in PBS) for 10 min at RT. After three subsequent washes in PBSA, embryos were transferred onto microscope slides and mounted using Fluoromount-G. Imaging was performed as described above.

### Embryo labelling quantification

Images were taken using a Leica DMI8 fluorescence microscope. Within each replicate, intensity and gain were identical to ensure differences in fluorescence intensity were not due to technical factors. Embryonic CDX2, 5meC, and Hoescht intensities were quantified using the *Measure* function within Fiji (ImageJ). The CDX2 or 5meC to Hoescht signal ratio for each embryo was calculated using Microsoft Excel.

### Statistical analyses

Additional statistical analyses not previously described were performed using GraphPad Prism for Mac (v. 10.4.2, b. 534). More specifically, organoid morphologies (**Fig**. **1F-H** & 3E-G) were compared by two-way analysis of variance (**ANOVA**) with multiple comparisons and subsequent Tukey’s *post hoc* test. Intra-organoid fluid relative volumes (**Fig**. **2F** & **4A**), embryo cleavage and blastocyst rates (**Fig**. **5C-D**), and relative fluorescent intensities (**Fig**. **5G-H**) were compared by ordinary one-way ANOVA, without matching or pairing, with multiple comparisons, and subsequent Tukey’s *post hoc* test. Asterisks denote significance as follows: *P*≤0.0001 (****); *P*≤0.001 (***); *P*≤0.01 (**); and *P*≤0.05 (*).

## ACKNOWLEDGEMENTS

The authors thank Mr. Manuel “Boo” A. Persica in the Louisiana State University (**LSU**) School of Animal Sciences and the staff at Coastal Plains Meat Company, Eunice, Louisiana, for their assistance with tissue collection. Figure 5I was created in part using biorender.com. The authors also acknowledge the use of ChatGPT for improving sentence structure and readability in parts; however, the authors take full responsibility for the content of this manuscript. This work was funded by: The LSU Agricultural Center Therapeutic Cannabis Research Committee (PG010126) to CAS (PI), XF (Co-I), and KRB (Co-I); The State of Louisiana Board of Regents [LEQSF(2023-26)-RD-A-03] to CAS (PI) and XF (Co-I); The United States Department of Agriculture (USDA) Research Capacity (Hatch) funds (LAB-94578) to CAS (PI); and The LSU Agricultural Center Collaborative Research Program (PG010315) to CAS (PI), XF (Co-I), and AV (Co-I). AV is funded in part by the State of Louisiana Board of Regents [LEQSF(2023-26)-RD-A-16].

## AUTHOR CONTRIBUTIONS

Conceptualization: DMF & CAS. Methodology: DMF, ID, CCW, AV, KRB & CAS. Investigation: DMF, ID, ZLB, YL, XZ & XF. Resources: XF, KRB & CAS. Data curation: DMF & CAS. Formal analysis: DMF, MFD, AV & CAS. Manuscript preparation: DMF, AM, ALP & CAS. Project supervision: CAS. Funding acquisition: CAS.

---

**Supplementary Table 1**. List of antibodies.

**Supplementary Table 2**. List of culture and other media compositions.

**Supplementary Table 3**. Total organoid RNA recovered.

**Supplementary Table 4**. Raw RNA-seq counts.

**Supplementary Table 5**. Raw metabolomic peak areas.

